## Supplementary material for "Development and Characterization of PAR-Trackers: New Tools for Detecting Poly(ADP-ribose) In Vitro and In Vivo": Figure 2 - Figure Supplement 2, video

### Slide 1
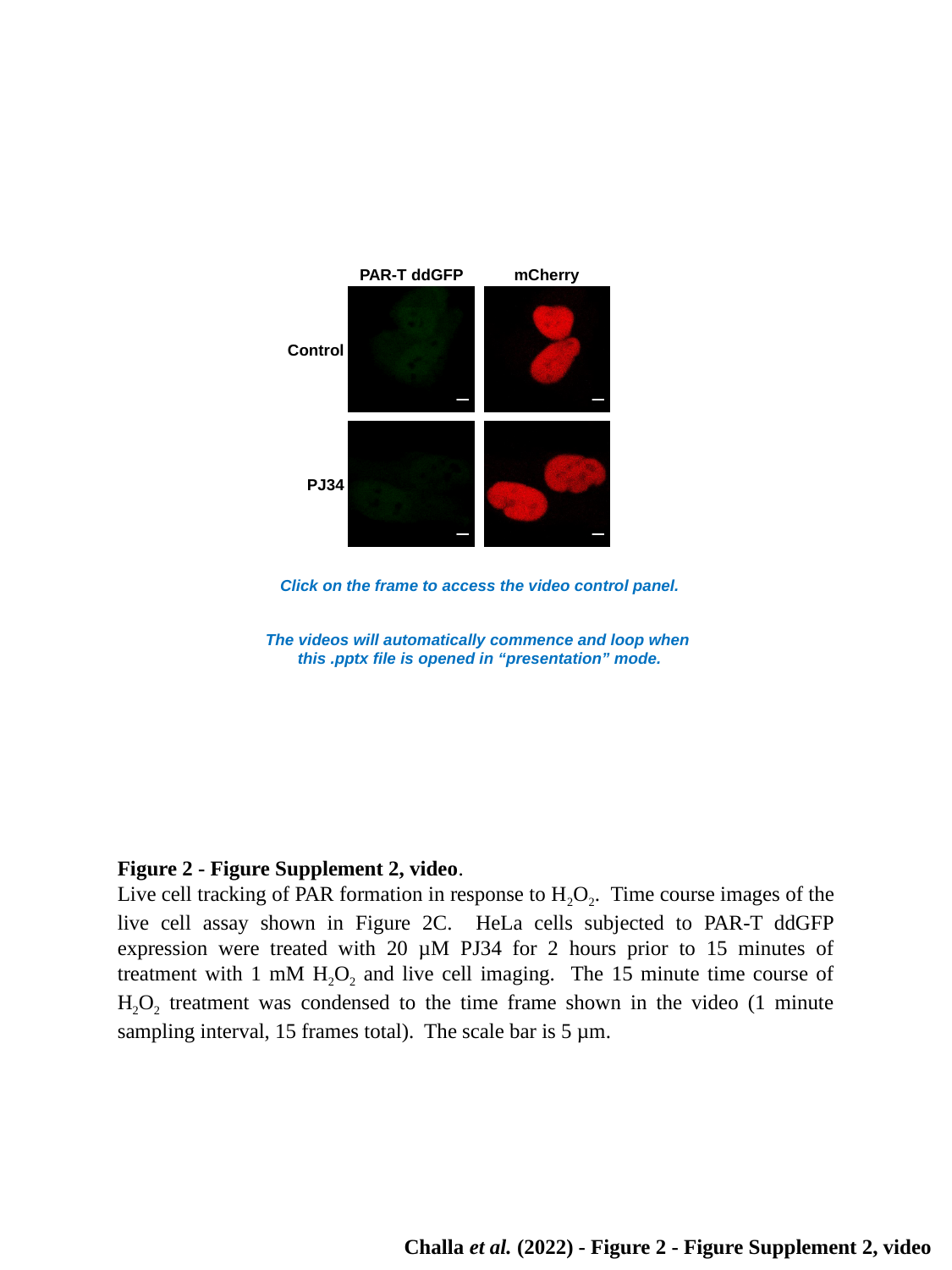

PAR-T ddGFP
mCherry
Control
PJ34
Click on the frame to access the video control panel.
The videos will automatically commence and loop when
this .pptx file is opened in “presentation” mode.
Figure 2 - Figure Supplement 2, video.
Live cell tracking of PAR formation in response to H_2_O_2_. Time course images of the live cell assay shown in Figure 2C. HeLa cells subjected to PAR‑T ddGFP expression were treated with 20 µM PJ34 for 2 hours prior to 15 minutes of treatment with 1 mM H_2_O_2_ and live cell imaging. The 15 minute time course of H_2_O_2_ treatment was condensed to the time frame shown in the video (1 minute sampling interval, 15 frames total). The scale bar is 5 µm.
Challa et al. (2022) - Figure 2 - Figure Supplement 2, video
